# Legacy effects of European colonialism on hotspots of biocultural diversity threat

**DOI:** 10.1101/2025.01.27.635019

**Authors:** Bernd Lenzner, Andreas Baumann, Sietze Norder, Franz Essl, Hannes A. Fellner

## Abstract

Patterns of biological diversity have been shaped by cultural practices in the past, while in turn, cultures and languages have evolved in close interaction with local species and ecosystems. However, in the Anthropocene, humans are putting increasingly more and diverse pressures on ecosystems and cultures resulting in accelerating threat levels on both. Understanding where biological and linguistic diversity (a common measure of cultural diversity) is threatened globally, and which drivers shape their distribution is crucial for pinpointing hotspots and prioritizing efforts to counter these threats. We find that the spatial patterns of the erosion of biological and cultural diversity are weakly congruent on a global scale and that they are driven by differential sets of mechanisms. However, our results also indicate that particularly large-scale historic socio-political events like the European colonial expansion have left long-lasting imprints on both biological and cultural diversity. The duration of European colonialism (i.e., the time a specific region was occupied by Europeans powers) significantly increases contemporary observed threat levels of biological and linguistic diversity.

**Significance Statement:** Biological and cultural diversity are closely intertwined. However, in an increasingly changing world where anthropogenic drivers put increasing pressures on ecosystems and cultures, both are threatened. Understanding how these threats to both, biological and linguistic diversity, are organized in space and where their individual and combined threat hotspots are is crucial for prioritization efforts to safeguard both. We also show what drivers affect their threat level globally and pinpoint that particularly large-scale socio-political events like the European colonial expansion have left long-lasting imprints on both biological and cultural diversity.

## Main Text Introduction

Nature and culture in the Anthropocene are in peril. Biological diversity is rapidly declining with extinction rates being 100-1000 times higher compared to pre-human times and an estimated one million species at risk of extinction (1–3). Simultaneously, language diversity, which often functions as a proxy of cultural diversity, also declines rapidly (4, 5). At least 40% of all languages are classified as threatened (6) and thousands of languages went extinct over the past 500 years (7, 8). The synchrony of these patterns is likely no coincidence but associated with the increasing alteration of the biotic and abiotic face of the earth by human activities, including climate change, resource extraction, overexploitation, pollution and land/sea-use change. In particular, cultural and socio-political dynamics - having been profoundly altered by colonialism, subsequent globalization and (inter)national policies - further affect nature and culture (1, 9–11).

Biological and linguistic diversity are intertwined at a global scale, showing congruencies in their spatial diversity patterns (4, 12–15). This biocultural diversity (i.e., the combination of biological, cultural diversity, where linguistic diversity often serves as a substitute for cultural diversity; see for example (13)) is particularly high in topographically and climatically complex regions in Mesoamerica, Western Africa, the Indo-Malay region and Melanesia (4, 13–15). Another distinctive characteristic of such hotspots is their association with lands of indigenous people and local communities that cover around 30% of the global land surface and at least 40% of protected areas (16, 17). Also, most threatened languages are spoken by indigenous and local communities (9, 18). While the loss of biodiversity threatens ecosystem functioning and human livelihoods globally(1), the loss of linguistic and cultural diversity erodes cultural identities, as well as unique knowledge systems, such as those related to medical use of plants for modern medicine (19, 20) or sustainable harvesting techniques (21).

While contemporary pressures pose important direct threats to biocultural diversity, historical dynamics can leave enduring legacies that remain detectable today. European colonial expansion, characterized by land appropriation, exploitation, and dramatic reorganization of populations through the slave trade and settler colonialism, led to large-scale exploitation of natural resources, landscape conversion, and habitat fragmentation. This resulted in the destruction of unique ecosystems, loss of native biodiversity, introduction of alien species, expropriation of indigenous lands, and the displacement, expulsion, and genocide of native peoples (1, 9, 11, 22, 23). In addition, other socio-political pressures resulted in the erosion and loss of linguistic and cultural diversity through colonial assimilation policies or the destruction of cultural and other heritage sites (24–26), as did colonially induced diseases (8, 27, 28). Lag effects are a common phenomenon in biocultural change (29–31). However, it remains unclear to which extent legacies of European colonialism continue to impact contemporary biocultural diversity and its associated threat globally. Here, we combine several global datasets on threat status of languages (using information from Ethnologue (32)) and species (using the global IUCN Red List assessments for amphibians, birds, mammals and reptiles (33)). Using the IUCN Red List Indicator (RLI) framework, we calculated three different threat indicators relating to the threat of (i) linguistic diversity (*RLI*_*linguistic*_), (ii) biological diversity (i.e., threat of amphibians, birds, mammals and reptiles; *RLI*_*biological*_), and (iii) biocultural diversity (i.e., mean threat of linguistic and biological diversity; *RLI*_*biocultural*_) for 176 countries. The RLI was developed as a measure of species threat in a specific region (34, 35) based on individual threat assessment for the species present in that region. It is bound between 0 and 1 where 0 indicates that all species in a region are extinct and 1 that none are threatened (see supplementary methods and table S1). Here, we use this conceptual framework to also assess the threat status of languages (as has been done before; (4, 36, 37)). Specifically, we analyse if hotspots of threatened linguistic and biological diversity are spatially congruent.

In a second step, we investigate the factors which explain these patterns using beta regression mixed-effect models. We particularly focus on testing if European colonial history has left lasting imprints on current threat levels of linguistic, biological and biocultural diversity. We also examine how strong this impact is in relation to contemporary variables related to environmental and anthropogenic change.

## Results and Discussion

### Hotspots of threatened biocultural diversity

Linguistic diversity is most threatened in Australia, parts of Mesoamerica (i.e., El Salvador, Nicaragua), Israel, and Taiwan, while threats to biological diversity are most prevalent in Hispaniola (i.e., the Dominican Republic, Haiti), and (sub)tropical island nations such as Madagascar, Mauritius, and New Zealand (figure 1 A & B; table S3). When comparing the two patterns we find only a weak positive correlation in the spatial distribution pattern of linguistic and biological diversity threat (Pearson correlation = 0.14; figure 2A & S7). Overall, biodiversity is more threatened in regions of tropical and central Asia, Central America, as well as Northern and Southeastern Africa (turquoise regions in figure 2B; table S3), whereas linguistic diversity is more threatened across North and South America, Northeastern and Southern Africa and Australia (purple regions in figure 2B). Looking at biocultural diversity (i.e., measured as the mean threat across linguistic and biological diversity) we see high threat levels particularly in Oceania (i.e., Australia, New Zealand) and East Asia (i.e., Japan, Taiwan), and Haiti (figure 1C; table S3).

**Figure 1.**
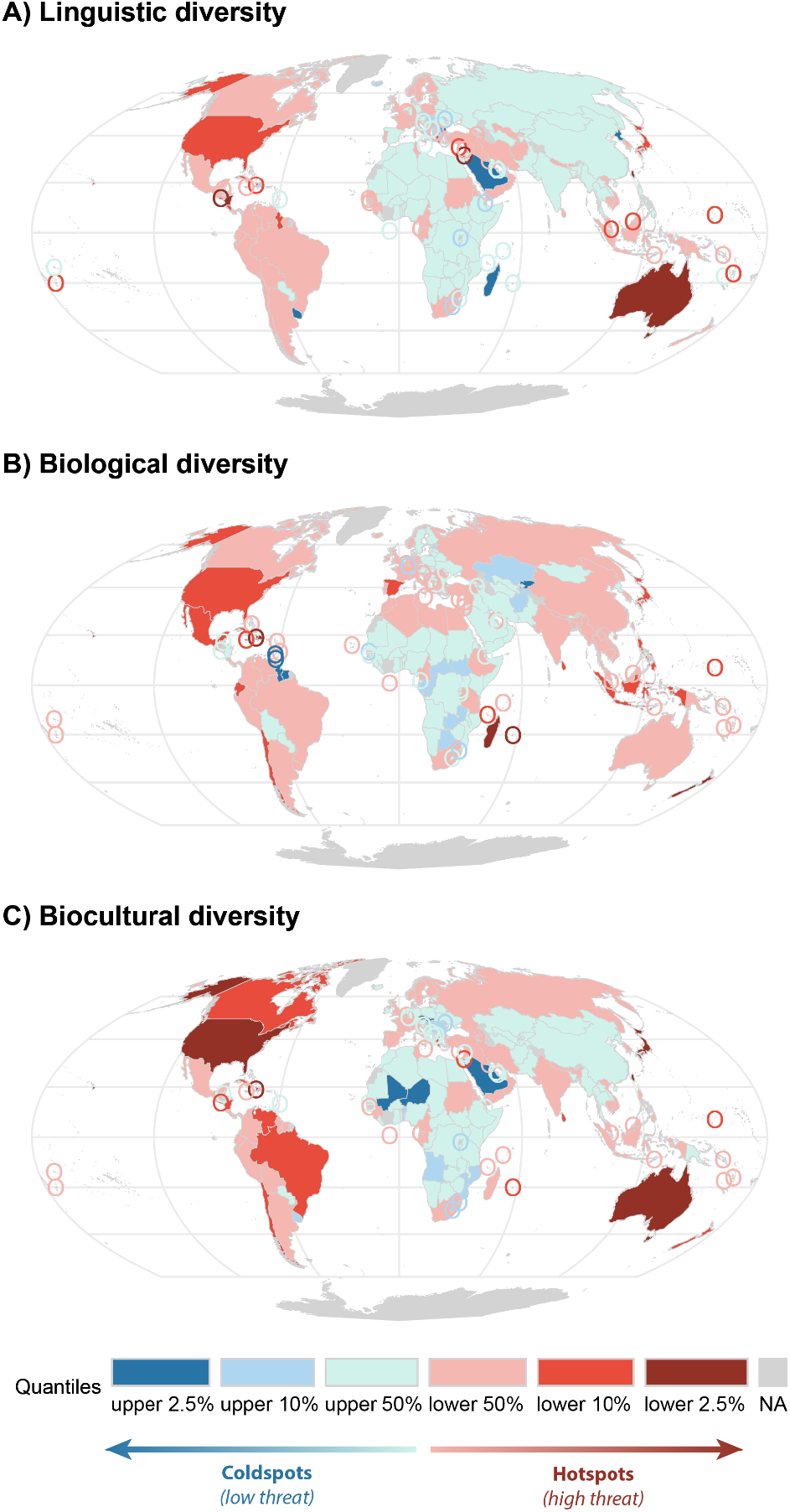
Global hot- and coldspots of the threat to (A) linguistic (RLIlinguistic), (B) biological (RLIbiological), and (C) biocultural diversity (RLIbiocultural). Hotspots are defined as regions in the lower 2.5% quantile corresponding to low values of RLI and coldspots are defined as regions in the upper 2.5% quantile, corresponding to high values of RLI. Small regions and islands are encircled using the same color coding.

**Figure 2.**
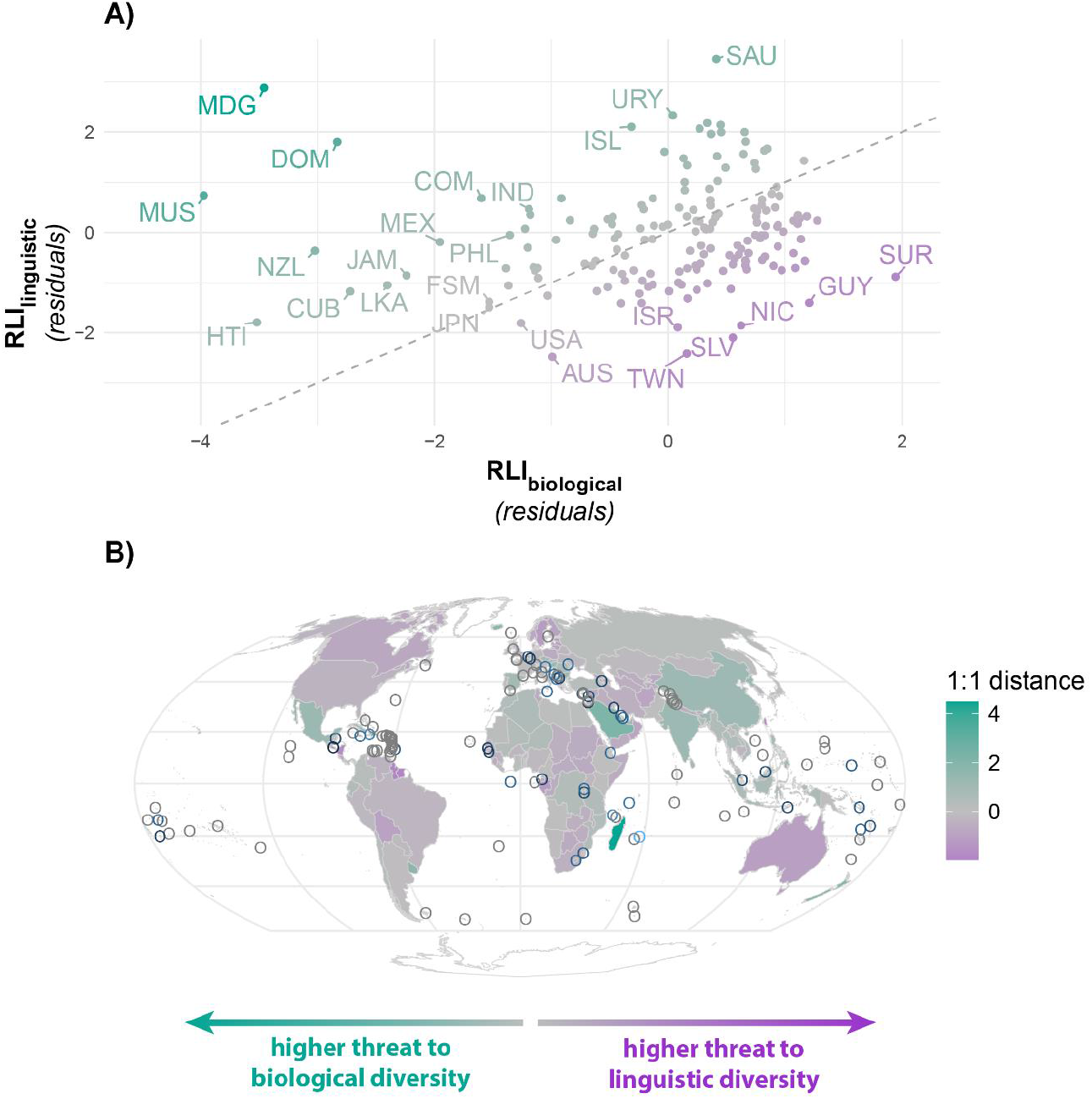
(A) Correlation (using the residuals after correcting for area and sampling bias) between the threat levels of linguistic (RLIlinguistic) and biological (RLIbiological) diversity based on the hotspot analysis and (B) its spatial distribution. The grey dashed line indicates the 1:1 line of perfect correlation between RLIlinguistic and RLIbiological. Values above the 1:1 line (in green) indicate countries with higher biological diversity threat levels compared to linguistic diversity threat levels and values below the 1:1 line (in purple) indicate the opposite pattern. Color intensity indicates distance to the 1:1 line with darker colors being more distant and lighter and grey colors closer to the line. Abbreviations follow the ISO-3 codes for countries.

### European colonialism left long-lasting imprints on threats to biocultural diversity

Among all predictors considered in this study, the time that a region was under European colonial rule (henceforth “occupation time”) is the only one that negatively affects both linguistic and biological diversity. In other words, the longer a region was occupied, the greater the current threats to its linguistic and biological diversity. Our results thus show that colonialism, as a disruptive process, has consistently impacted both biological and linguistic diversity, though likely through different mechanisms (13, 14, 38). Unsurprisingly, we see the same relationship in the compound metric on biocultural diversity.

The rise of European colonialism brought about economic and societal changes that directly impacted biodiversity. For example, the introduction of new crops, expansion of agricultural areas, development of infrastructure, overhunting, and the spread of invasive alien species all affected local ecosystems (39, 40). In extraction colonies, where the focus was on producing raw materials (like cotton and sugar in Mozambique or rubber in Southeast Asia (41)), large-scale land conversion led to the destruction of natural ecosystems and their species (39, 42). Similarly, managerial and mission colonies, which drove natural resource exploitation and the spread of religious beliefs, also contributed to species decline through practices like hunting for fur or wild meat, leading to significant reductions in species populations and even local or global extinctions (39, 42, 43).

Additionally, European colonialism led to the introduction of alien species for various reasons and purposes (22). Some of these species had harmful effects on native flora and fauna, especially on islands (44, 45). The legacy of this species redistribution during colonial times is still evident today in the composition of regional alien species pools (23, 45). Finally, we see differences in the patterns across taxonomic groups (Table S6). In our results, particularly groups of larger bodied animals that were exploited for their meat, fur or ornamental purpose like birds, mammals and reptiles, show imprints of European colonialism. Amphibians on the other hand, do not show such a significant legacy effect, likely given that amphibians exerted lower levels of direct exploitation and current threat patterns are driven more strongly by contemporary threats like land conversion or alien pathogens like the amphibian chytrid fungi *Batrachochytrium dendrobatidis* and *B. salamandrivorans* (46).

Similarly, European colonialism had severe impacts on linguistic diversity (11). While different European empires imposed different impacts on local languages associated with the style of occupation and established empire policies (11, 47, 48), some commonalities have been observed across empires. The introduction of European diseases, violent conflicts with local communities, and the forceful occupation of lands led to the loss of local languages and cultures, as well as the expulsion and forced relocation of people from their lands. This led to a dispersion of speakers of the same mother tongue where the local language got lost as vernacular (11). In previously isolated regions, the introduction of novel diseases caused the collapse of human populations, and, consequently, the loss of their languages. Additionally, in regions with many native languages spoken and in settler colonies dominated by colonialists of European descent, the imperial language was established as lingua-franca of commerce and administration. Sometimes, this resulted in the formation of pidgins and creoles (49). Regardless, integration of local populations into European religious, social, and economic systems had people both willingly and unwillingly to abandon their own language(s) in favor of a language that was more prestigious and from a communicative perspective more convenient in the newly established socio-economic environment (8, 50–52).

### Contemporary anthropogenic pressures continue to increase biocultural threat levels

Current human activities continue to erode both biological and cultural diversity worldwide (IPBES 2019; Bromham et al. 2022). We observe that higher levels of anthropogenic pressures, such as urbanization, intensive agriculture, and per capita GDP as an integrative measure of economic activity (Figure 3, Table S5), are strongly associated with increased threats to both types of diversity reflecting the impact of intensified trade, urban growth, and land transformation. In addition, smaller regions exhibit a significantly higher risk of losing linguistic diversity. Similarly, for biological diversity, roughness significantly increases threat levels (Figure 3, Table S5).

**Figure 3.**
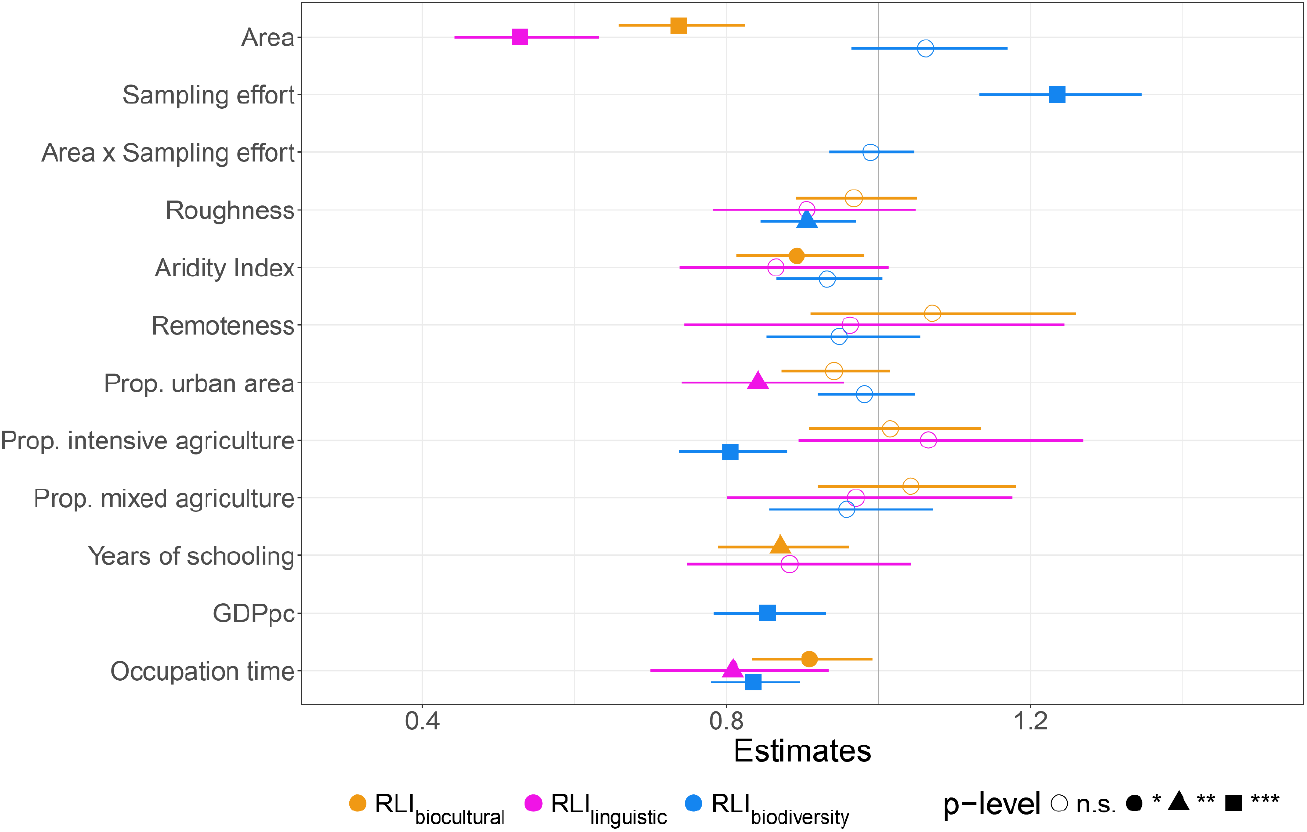
Coefficients from the beta regression models analyzing the relationship between the three different diversity threat measures (orange: RLIbiocultural; pink: RLIlinguistic, blue: RLIbiodiversity) and the associated predictor variables. Significant levels relate to the following values: n.s.: p > 0.01; *: p < 0.01; **: p < 0.05; *** p < 0.001.

We found that roughness is associated with higher threat to biodiversity. While many mountains are less used by humans(53), they harbor species that often have narrow range sizes and subsequently a higher likelihood of becoming threatened or locally extinct (54). A similar mechanism has been proposed for linguistic diversity, where high diversity has been observed in topographically complex regions like New Guinea (55), however also topographically less complex regions, like wetlands, have been important to humans as they provide valuable resources (56, 57). Such dynamics might explain why we do not find congruent patterns between linguistic and biological diversity. The absence of an effect of education on language endangerment (e.g., found in (10)) might be a result of the spatial scale of the analysis and the absence of data on education and years of schooling, especially data if education supports or restricts the learning of minority languages are missing.

### Understanding threats to biocultural diversity needs to incorporate a historical perspective

Despite the vastly different timescales over which patterns of biodiversity and cultural diversity have evolved, current threat levels of both species and languages are deeply rooted in the legacies of European colonial expansion. Our study is the first to demonstrate that the threats to biological and linguistic diversity are deeply interconnected on a global scale, and that the European expansion was the dominating force that – for the first time in human history – created a truly globally connected world. Even decades after the collapse of the European empires, their legacy continues to influence current threat levels of both languages and species. Understanding and learning from such past socio-political events and their long-term consequences on biological and linguistic diversity is essential for addressing and alleviating contemporary pressures.

## Materials and Methods

### Measuring language and biodiversity threat

To measure biocultural diversity threat (i.e., encompassing biological, cultural and linguistic diversity) we used the well-established Red List Indicator of biological diversity threat (RLI_biological_; (34, 58)) and built an equivalent for linguistic diversity threat (RLI_linguistic_) by mapping the EGIDS language endangerment classification scheme onto the Red List threat assessment categories (see (4, 36, 55) and supplementary material). The RLI_biological_ uses data from the IUCN Red List of Threatened Species (IUCN Red List, http://www.iucnredlist.org), widely recognized as the most authoritative, objective and comprehensive approach for evaluating the global conservation status of species and categorizing them according to their risk of extinction (34, 35, 58, 59). Here we focus on well-sampled taxonomic groups to quantify biodiversity (namely amphibians, birds, mammals, and reptiles). Both indicators the RLI_biological_ and RLI_linguistic_ quantify the mean threat level for biological and linguistic diversity at country level and are bound between 0 (high threat, meaning all languages/species listed as extinct) and 1 (no threat, meaning all languages/species are listed as least concern).

Analyses were performed for six different diversity threat measures: (i) RLI_biocultural_ = biocultural diversity (i.e., calculated as the mean of the RLI_biological_ and RLI_linguistic_), (ii) RLI_linguistic_ = linguistic diversity, (iii) RLI_biological_ = biological diversity (i.e., including data of all 4 taxonomic groups), (iv) RLI_amphibians_, (v) RLI_birds_, (vi) RLI_mammals_ and (vii) RLI_reptiles_ (the latter four are presented in supplementary material).

### Hotspot analysis

To understand the spatial patterns of the seven diversity metrics we identified hot- and coldspots for each one separately. In a first step, we accounted for the difference in country size among regions by running generalized linear models with a beta error distribution (hereafter beta regression models) to account for response variables bound between 0 and 1. The models were fitted within the Generalized Additive Models for Location, Scale and Shape (GAMLSS) framework in R using the *gamlss* package (60). For RLI_biological,_ RLI_amphibians_, RLI_birds_, RLI_mammals_, and RLI_reptiles_ we additionally accounted for sampling effort (see below). Both predictors were log-transformed and scaled to increase model fit and ensure comparability between the predictors. Models were checked for spatial autocorrelation visually using acf-Plots as well as Durbin-Watson test (61).

To account for heterogeneous knowledge of taxonomic data worldwide we used a proxy for sampling effort, based on global occurrence records for well-known vertebrate species (amphibians, mammals, and birds) based on data from (62). The proxy was calculated by estimating the percentage of completeness of species occurrences stored in GBIF at the grid cell level. We then aggregated this information to the country level to obtain a mean sampling effort value (following the procedure in (63)). Sampling effort was then included as a separate and an interaction term with country area in the models. Unfortunately, such an indicator of sampling effort is, to our knowledge, not available for linguistic diversity. We also do not think that the sampling bias proxy for biodiversity is transferable to linguistic diversity data globally, given that linguistic data sampling within the Ethnologue had a strong historical focus on regions outside of Europe and Northern America, contrary to the observed sampling bias in biological diversity. Hence, we corrected linguistic diversity for area of the country only.

In a final step we used the unexplained variance (i.e., deviance residuals) at country level from the models to identify hot- and coldspots across diversity metrics. For that the residuals were categorized into quantiles (2.5%, 10%, 50%, 90% and 97.5%). Regions in the lower 2.5% quantile are defined as hotspots of diversity threat (i.e., low values of RLI) and regions in the upper 2.5% are defined as coldspots of diversity threat (i.e., high values of RLI).

### Analysis of drivers of diversity threat

To identify the drivers of threat levels across diversity measures, we build seven separate beta regression models for the respective diversity threat metrics as response variables and a set of predictors tailored to the different diversity threat metrics. Predictors included geographic (i.e., area, habitat heterogeneity), climatic (i.e., aridity), socio-economic (i.e., remoteness, proportion of intensive agricultural land, proportion of mixed agricultural land, proportion of urban areas, level of education, per capita gross domestic product), socio-political (i.e., occupation time by the European empires) and sampling effort (for all biological diversity related indicators). For a detailed description of all variables see supplementary material.

The six separate models again were built using a beta error distribution to test the relationships between the respective diversity threat indicators and the predictor variables. Each model included all geographic, climatic, socio-economic and socio-political variables, while the biological diversity related indicators models additionally included the sampling-effort term. For each model we also included a random effect for the region separating the globe into 17 geopolitical regions to account for spatial autocorrelation. Two variables, GDPpc and Years of schooling were highly correlated (Pearson’s correlation coefficient >0.7; figures S5 and S6) and could not be used in the same model. Given that years of schooling has been identified as a potentially important driver of linguistic diversity threat (10), we decided to keep this variable for the models on biocultural and linguistic diversity. For biological diversity we kept GDPpc in the models as GDPpc used as a proxy for human influence on ecosystems as well as trade and infrastructure development has been shown to be one of the major drivers of biodiversity loss (1). We also ran the models for linguistic and biocultural diversity using GDPpc instead of years of schooling and results are reported in table S7. Severely right-skewed variables (e.g., proportion of intensive agricultural land, proportion of mixed agricultural land, proportion of urban areas, remoteness, and GDPpc) were log-transformed to ensure normally distributed residuals and all variables were centered and scaled to ensure comparability. Finally, predictor collinearity was tested using Pearson’s pairwise correlation to ensure that variable correlation was > -0.7 and < 0.7 ((64); see figures S5 & S6) and model fit was investigated visually to ensure variance homogeneity and normality of residuals (see figure S8). Residual autocorrelation in all models was tested using the Durbin-Watson test implemented through the *check_autocorrelation()* function in the R-package *performance* (65). None of the models showed any residual autocorrelation (i.e., all tests remained insignificant; p > 0.05).

