## Supplementary Information for "Legacy effects of European colonialism on hotspots of biocultural diversity threat"

‡ both authors should be considered last authors

*** corresponding author:** Bernd Lenzner

### Supporting Information Text

#### Data preparation

*Species and language data*

Native species data for amphibians, mammals, and reptiles were obtained from the IUCN Red List data portal (1). Bird data were obtained from Bird Life International (2). All datasets contained shapefiles of native species ranges for the respective taxonomic groups including the global threat status based on the Red List classification for each species. For birds we used a subset of species range polygons, excluding those indicating that the presence of the species is uncertain, where the species origin is listed as introduced, vagrant, uncertain or assisted colonization, and for species known in this region as migratory species or with uncertain seasonal occurrence, to keep the selection of species in line with the IUCN Red List data. After data cleaning we intersected all remaining range polygons with the administrative delineations at the country level (level 0) from the Database of Global Administrative Areas (GADM 2024, version 4.1) to obtain species country lists.

For language distributions, we used data provided by the Ethnologue (3), the currently most up-to-date and complete repository of the languages of the world. The Ethnologue database export that we used consists of information for 7020 languages. We decided to keep our selection of languages as comprehensive as possible and kept all languages in Ethnologue for which information about allocation to countries (ISO-2 codes) and their threat level (EGIDS, see below) was provided. This includes pidgin languages, creole languages, languages that have emerged through language contact in the past (4), as well as sign languages. Since, due to historical migration events, it is not trivial to decide, which languages count as 'native' in analogy with the species data (and since there exists, to the best of our knowledge, no comprehensive dataset providing such a classification), no specific language family was excluded for any of the countries. This also means that we included colonial languages (and languages that have emerged from them) in our analysis. Note, though, that the fraction of colonial (here: European) languages and their derivates in colonized countries is typically very small, anyway. This resulted in complete data for 6921 distinct languages.

*Calculation of national threat levels*

National threat levels were calculated following the IUCN Red List Index framework for species threat. The Red List Index for biological diversity ($RLI_{biological}$) is calculated based on the global threat status assessment of species into five categories (LC = least concern, NT = near threatened, VU = vulnerable, EN = endangered, CR = critically endangered, and EX = extinct & extinct in the wild) following a distinct set of criteria (5, 6) using the formula:

$$RLI_{biological} = 1 - \frac{\sum_{s=1}^{N} W_{c,s}}{W_{EX} * N}$$

where N is the number of species, s refers to the individual species per region. $W_c$ is the weight of the threat category of the respective species and $W_{EX}$ is the weight of the threat category "extinct". Weights are assigned as follows: $W_{LC} = 0$, $W_{NT} = 1$, $W_{VU} = 2$, $W_{EN} = 3$, $W_{CR} = 4$, and $W_{EX} = 5$. The RLI is bound between 0 and 1, where 1 indicates the lowest threat level (i.e., all species listed as LC) and 0 indicates highest threat level (i.e., all species listed as EX).

We applied the same framework to estimate national threat levels of linguistic diversity to get a Red List Index of linguistic diversity ($RLI_{linguistic}$). To assign the IUCN threat categories to languages, we related them to the Ethnologue's Expanded Graded Intergenerational Disruption Scale (EGIDS) based on their provided definitions. Like the IUCN categories, EGIDS provides an estimate on language endangerment based on a set of criteria (e.g., including number of speakers, speaker population trends, speaker demographics, institutional status; SIL International 2022). A similar approach has been taken already in earlier studies (7–9). The relationships between the two schemes as used in our study are shown in table S1.

Finally, to get an estimate on the national threat level of biocultural diversity (i.e., the combination of linguistic and biological diversity; $RLI_{biocultural}$) we calculated the mean of both linguistic and biological diversity threat nationally.

**Table S1.** Description and relationship between the threat categories provided by the IUCN Red List and the Ethnologue EGIDS scheme.

| Red List Category | RedList description | EGIDS categories | EGIDS description | Threat category weight |
|---|---|---|---|---|
| **Least concern (LC)** | A taxon is Least Concern when it has been evaluated against the criteria and does not qualify for Critically Endangered, Endangered, Vulnerable or Near Threatened. Widespread and abundant taxa are included in this category. | **International** | The language is widely used between nations in trade, knowledge exchange, and international policy. | **0** |
| | | **National** | The language is used in education, work, mass media, and government at the national level. | |
| | | **Provincial** | The language is used in education, work, mass media, and government within major administrative subdivisions of a nation. | |
| | | **Wider Communication** | The language is used in work and mass media without official status to transcend language differences across a region. | |
| | | **Educational** | The language is in vigorous use, with standardization and literature being sustained through a widespread system of institutionally supported education. | |

| | | | | |
|---|---|---|---|---|
| **Near threatened (NT)** | A taxon is Near Threatened when it has been evaluated against the criteria but does not qualify for Critically Endangered, Endangered or Vulnerable now, but is close to qualifying for or is likely to qualify for a threatened category in the near future | **Developing** | The language is in vigorous use, with literature in a standardized form being used by some though this is not yet widespread or sustainable. | **1** |
| | | **Vigorous** | The language is used for face-to-face communication by all generations and the situation is sustainable. | |
| **Vulnerable (VU)** | A taxon is Vulnerable when the best available evidence indicates that it meets any of the criteria A to E for Vulnerable (see Section V), and it is therefore considered to be facing a high risk of extinction in the wild | **Threatened** | The language is used for face-to-face communication within all generations, but it is losing users. | **2** |
| | - Reduction in population size (approx. >30%) | | | |
| | - Geographic range severely fragmented, continuously declining, strongly fluctuating | | | |
| | - Population very small or restricted to a small area | | | |

| | | | | |
|---|---|---|---|---|
| **Endangered (EN)** | A taxon is Endangered when the best available evidence indicates that it meets any of the criteria A to E for Endangered (see Section V), and it is therefore considered to be facing a very high risk of extinction in the wild | **Shifting** | The child-bearing generation can use the language among themselves, but it is not being transmitted to children. | **3** |
| | | **Moribund** | The only remaining active users of the language are members of the grandparent generation and older. | |
| **Critically Endangered (CR)** | A taxon is Critically Endangered when the best available evidence indicates that it meets any of the criteria A to E for Critically Endangered (see Section V), and it is therefore considered to be facing an extremely high risk of extinction in the wild. | **Nearly Extinct** | The only remaining users of the language are members of the grandparent generation or older who have little opportunity to use the language. | **4** |
| **Extinct in the Wild (EX)** | A taxon is Extinct in the Wild when it is known only to survive in cultivation, in captivity or as a naturalized population (or populations) well outside the past range. A taxon is presumed Extinct in the Wild when exhaustive surveys in known and/or expected habitat, at appropriate times (diurnal, seasonal, annual), throughout its historic range have failed to record an individual. Surveys should be over a time frame appropriate to the taxon's life cycle and life form. | **Dormant** | The language serves as a reminder of heritage identity for an ethnic community, but no one has more than symbolic proficiency. | **5** |

| | | | |
|---|---|---|---|
| **Extinct (EX)** | A taxon is Extinct when there is no reasonable doubt that the last individual has died. A taxon is presumed Extinct when exhaustive surveys in known and/or expected habitat, at appropriate times (diurnal, seasonal, annual), throughout its historic range have failed to record an individual. Surveys should be over a time frame appropriate to the taxon's life cycle and life form. | **Extinct** | The language is no longer used and no one retains a sense of ethnic identity associated with the language. |

### Predictor data

*Habitat heterogeneity*

Habitat heterogeneity was calculated as the standard deviation of topographic complexity for each country based on the Shuttle Radar Topography Mission (SRTM) digital elevation model (DEM) at 30m resolution (10).

*Aridity*

The Global Aridity Index provides information on the available precipitation in an area as a function of precipitation, temperature and evapotranspiration. It is calculated based on global data for the years 1970 – 2000 at a 30' resolution and is calculated as:

$$Aridity\ Index = \frac{MA - Pr}{MA - ET_0},$$

with $MA - PR :=$ mean annual precipitation and $MA - ET_0 :=$ mean annual reference evapo-transpiration (11). Aridity index values can be categorized into five classes: <0.3 = hyper arid; 0.03 – 0.2 = arid; 0.2 – 0.5 = semi-arid; 0.5 – 0.65 = dry sub-humid; >0.65 humid. We calculated mean aridity index values for each country.

*Remoteness*

Remoteness here is defined as mean travel time to cities with a minimum population size of 5,000 and a maximum population size of 10,000 people based on (12). The dataset includes distances to 45,795 settlements as of the year 2015 with a combined sum of population of 322,797,326 inhabitants (12). Data is provided at 30 arc-second resolution. Distance to human settlements has been shown as a relevant proxy of linguistic and biodiversity, with more remote regions being more diverse (13, 14). Nelson et al. (12) provide different measures of travel times based on the size threshold for the considered cities. We decided for the distance to the smallest settlements, as those already can exert substantial pressure on linguistic and biodiversity. We also tested for correlation among the different measures and found a very high correlation between them (Pearson's r > 0.93; figure S1).

*Land cover variables and urbanisation*

For the proportion of agricultural land, we used the dataset provided by (15) who provide spatially explicit data on different food systems and other land cover classes, so called foodscapes. Specifically, we used data on (i) irrigated and/or intensive food production (hereafter called intense agriculture), (ii) mixed and diverse food cultivation (hereafter called mixed agriculture), and (iii) urbanized areas. Data are provided at a 5km spatial resolution. We calculated the mean area across all gird cells per country and divided it by the total area of the country. Proportion of agricultural areas and urbanisation acts as a proxy for anthropogenic impact on the environment resulting in land-cover change and increased anthropogenic pressure from natural to highly transformed landscapes.

*Level of education*

Data on the level of education was extracted from (16) available from the Wittgenstein Centre Data Explorer through the *wcde* – R package. Level of education is defined as the mean number of years a person spent in school averaged over the population cohort from the age of 20 to 64. Other measures for the level of education were available looking at different age cohorts, however, all were very highly correlation (Pearson's r > 0.83; figure S1) so we decided to use the broadest cohort.

Level of education was shown to be a correlate of linguistic diversity where higher education is associated with greater language endangerment as a consequence of for example reduced schooling of minority languages or the dominance of teaching a lingua-franca in the absence of minority education policies. On the other hand, policies to safeguard and support the learning of marginalized local and indigenous languages can result in a reduced endangerment of a specific language (14).

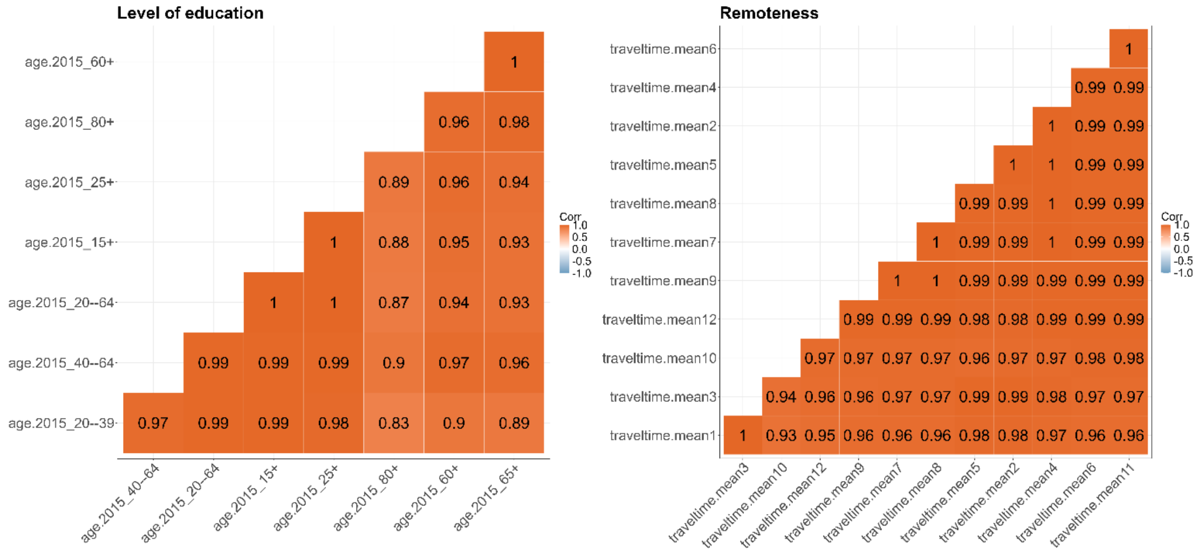

**Fig. S1.** Pearson's correlation between different measures for (A) level of education and (B) remoteness.

*GDPpc*

Gridded data on per capita gross domestic product (GDPpc) were obtained from (17). We used the data from the year 2015 at 5 arc-min resolution and calculated mean GDPpc values across countries. GDPpc is used as an indicator of human pressure, encapsulating multiple interconnected process including among others socio-economic development, trade, and urbanization. It has been shown that GDPpc negatively affects biodiversity and linguistic diversity (18).

*Occupation time*

Information on the occupation time was extracted from the ColDAT database (19). ColDAT is a compilation and harmonization of four individual datasets on the European empires. We used information for seven European empires: British, Spanish, Portuguese, Dutch, Belgian, German, and Italian empire. Occupation time was calculate based on the mean starting date and the maximum end date provided in ColDAT across all empires and is given in years. We used mean starting dates as they provide the earliest date of European colonial occupation provided in ColDAT often coinciding with the timing of formal integration of a colony into the empire. However, we note that colonial occupation started earlier than the formal integration with the establishment of trading posts, forts or other infrastructure related to managerial and mission colonies (20). Such information is difficult to obtain and using the mean starting date provides a conservative approach, rather underestimating the occupation time of a region and hence still providing robust results. Colonial occupation has been identified as one driver of linguistic and biodiversity loss (14, 20–23), however, previous publications have used this information generally as a binary variable (i.e., a region was occupied or not).

**Hotspot analysis**

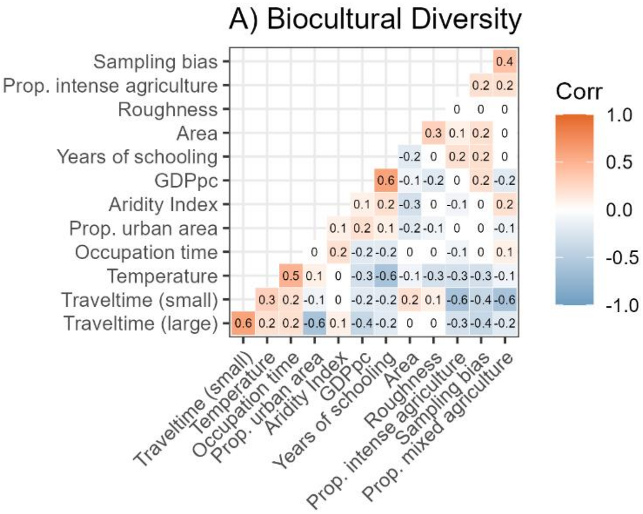

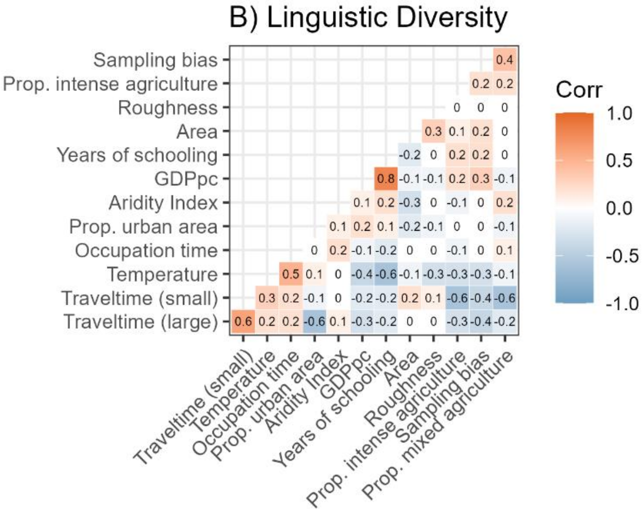

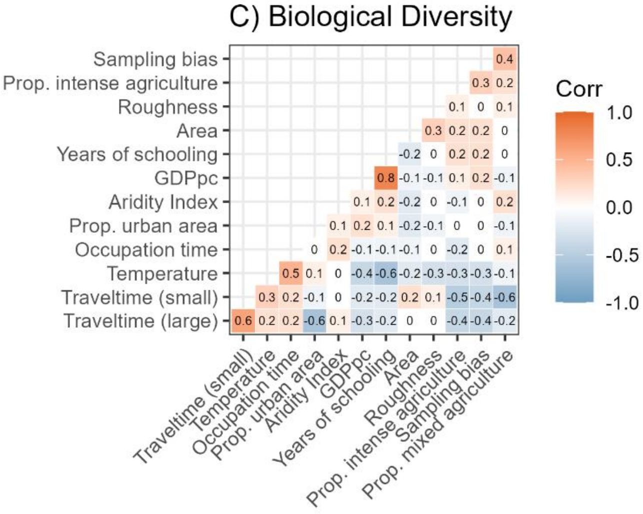

**Fig. S2.** Variable correlation of all potential driver variables for the different datasets used to explain patterns of threat to (A) biocultural diversity, (B) linguistic diversity, and (C) biological diversity.

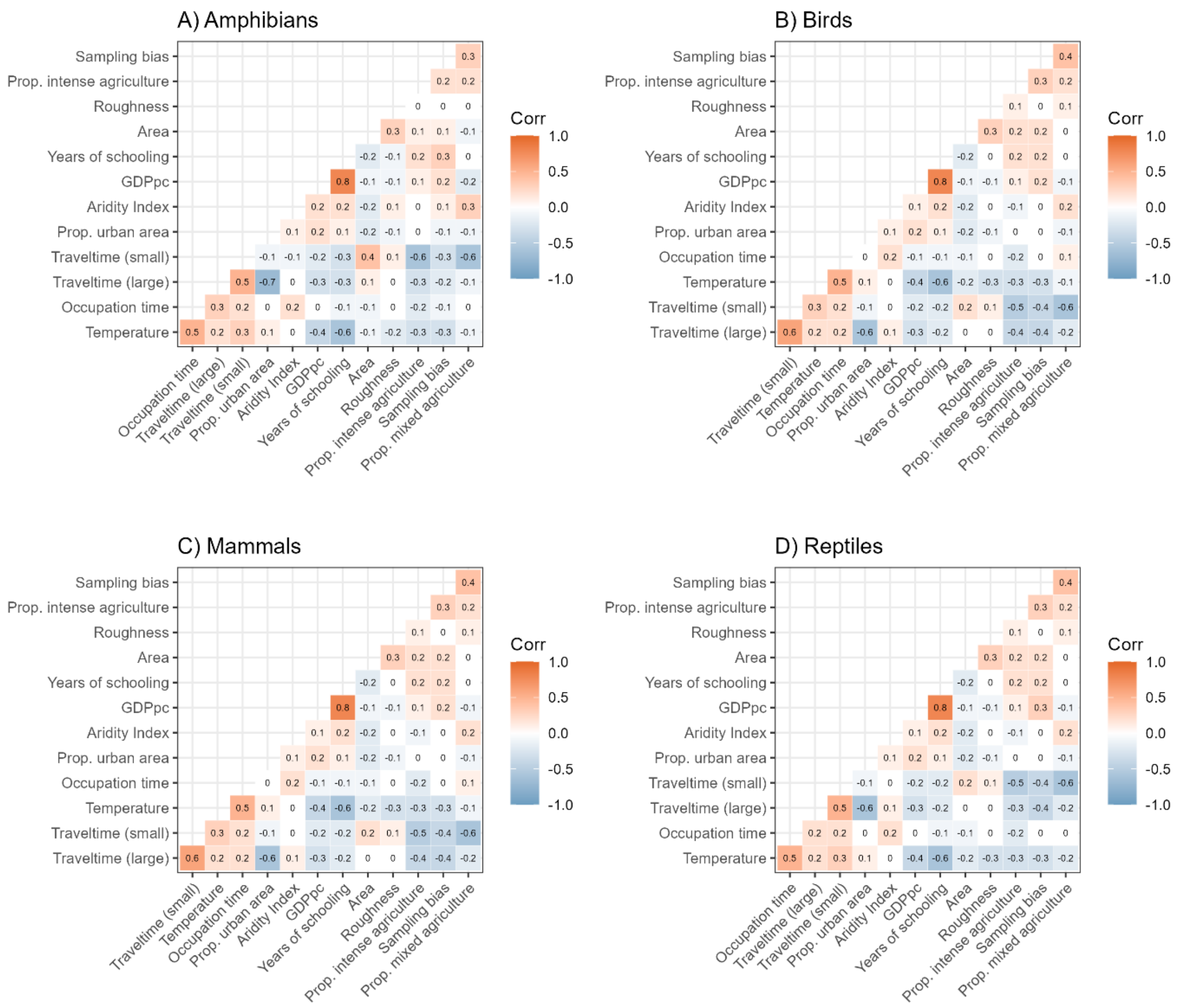

**Fig. S3.** Variable correlation of all potential driver variables for the different datasets used to explain patterns of biological diversity threat to (A) amphibians, (B) birds, and (C) mammals, and (D) reptiles.

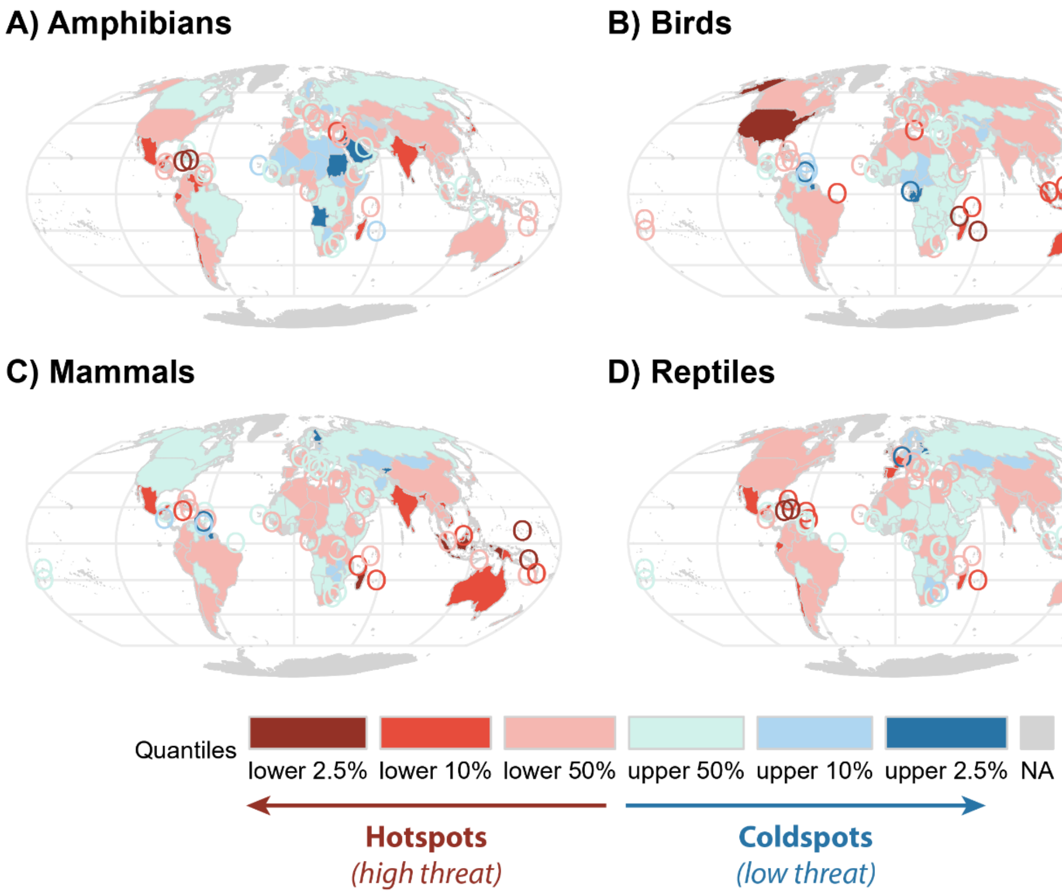

**Fig. S4.** Hotspots of biological diversity threat to (A) amphibians, (B) birds, (C) mammals, and (D) reptiles.

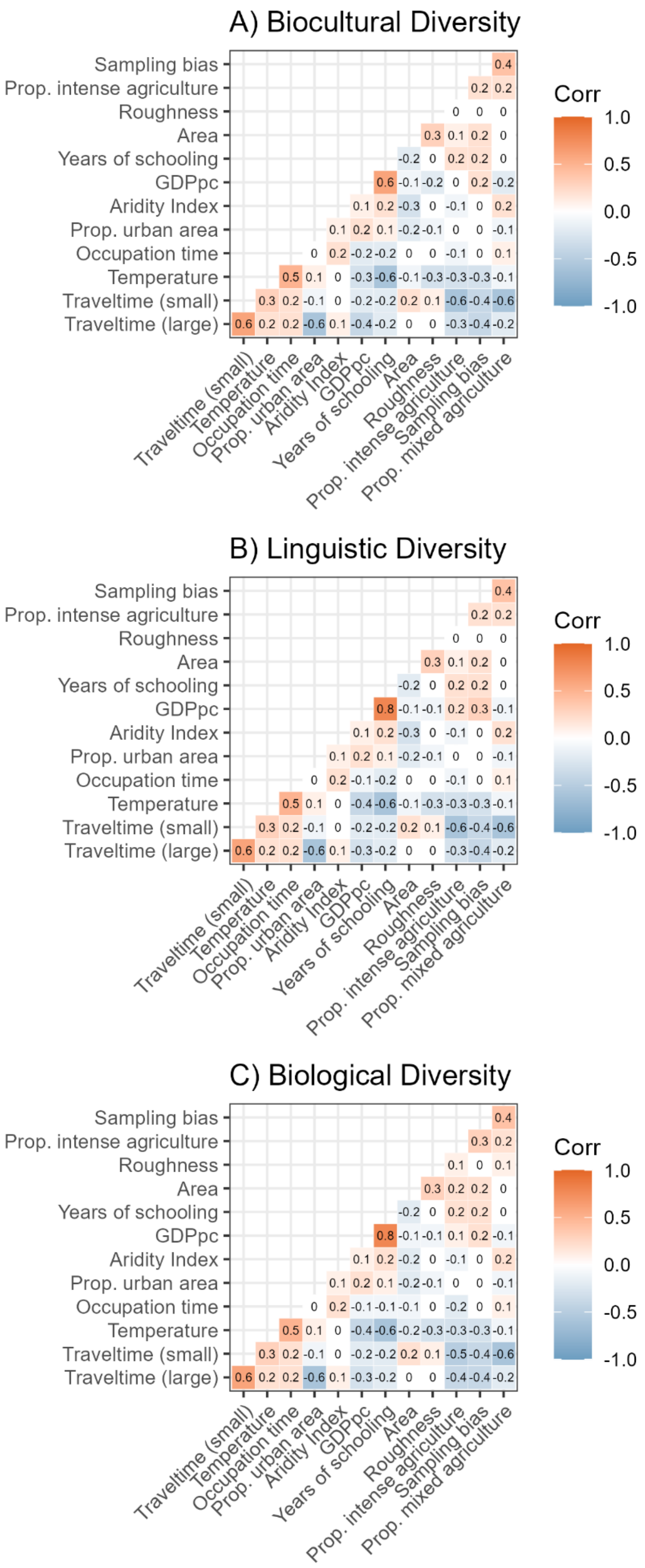
A) Biocultural Diversity
B) Linguistic Diversity
C) Biological Diversity
Sampling bias
Prop. intense agriculture
Roughness
Area
Years of schooling
GDPpc
Aridity Index
Prop. urban area
Occupation time
Temperature
Traveltime (small)
Traveltime (large)
Prop. mixed agriculture
Corr
1.0
0.5
0.0
-0.5
-1.0

**Fig. S5.** Pearson correlation of variables included in the driver models for (A) biocultural, (B) linguistic, and (C) biological diversity threat.

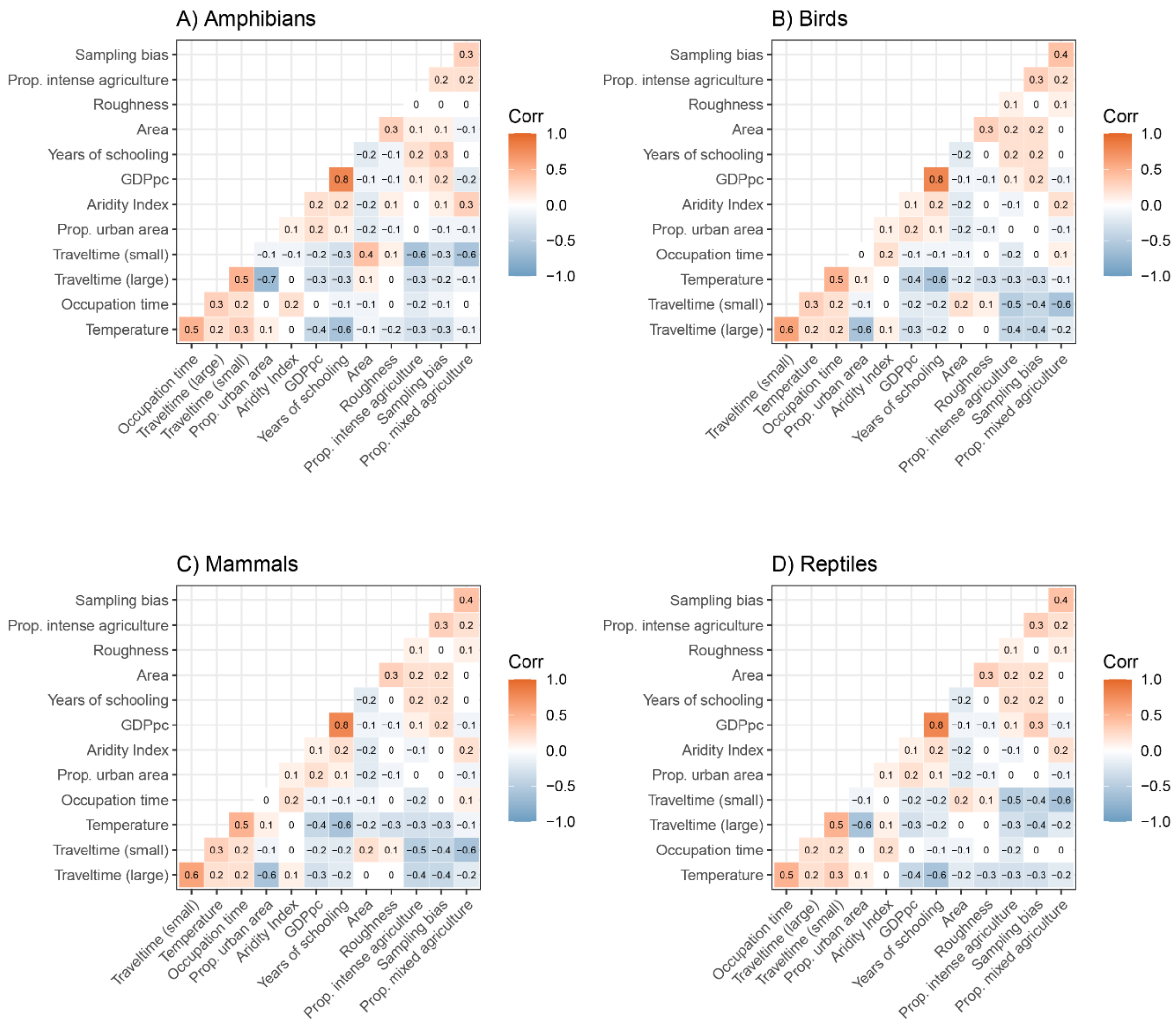

**Fig. S6.** Pearson correlation of variables included in the driver models for (A) amphibian, (B) bird, (C) mammal, and (D) reptile diversity threat.

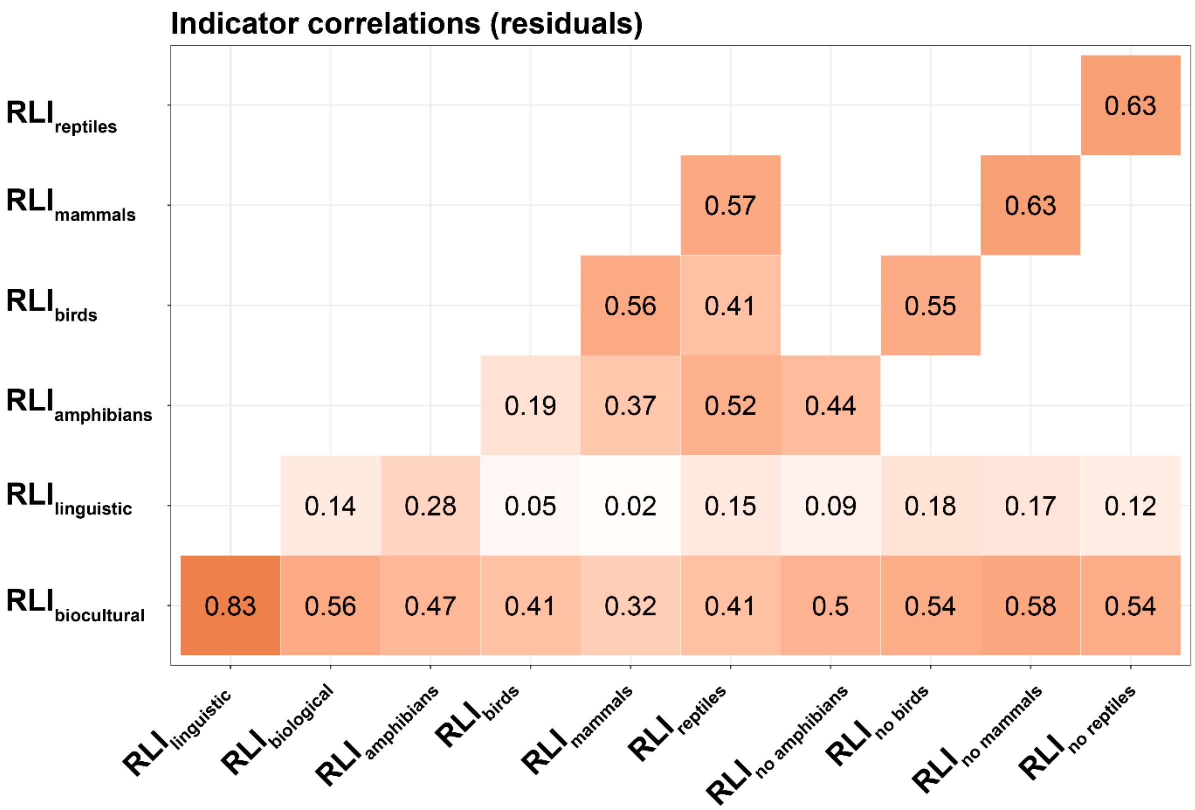

**Fig. S7.** Pearson correlation between diversity threat indicators. Correlation between $RLI_{biological}$ and the individual taxonomic groups (i.e., amphibians, birds, mammals, reptiles) was done by excluding the respective taxonomic group from $RLI_{biological}$ (i.e., $RLI_{no\ amphibians}$ represents the RLI per country including birds, mammals, and reptiles). RLI values are residuals after correcting for area and sampling bias (see methods section).

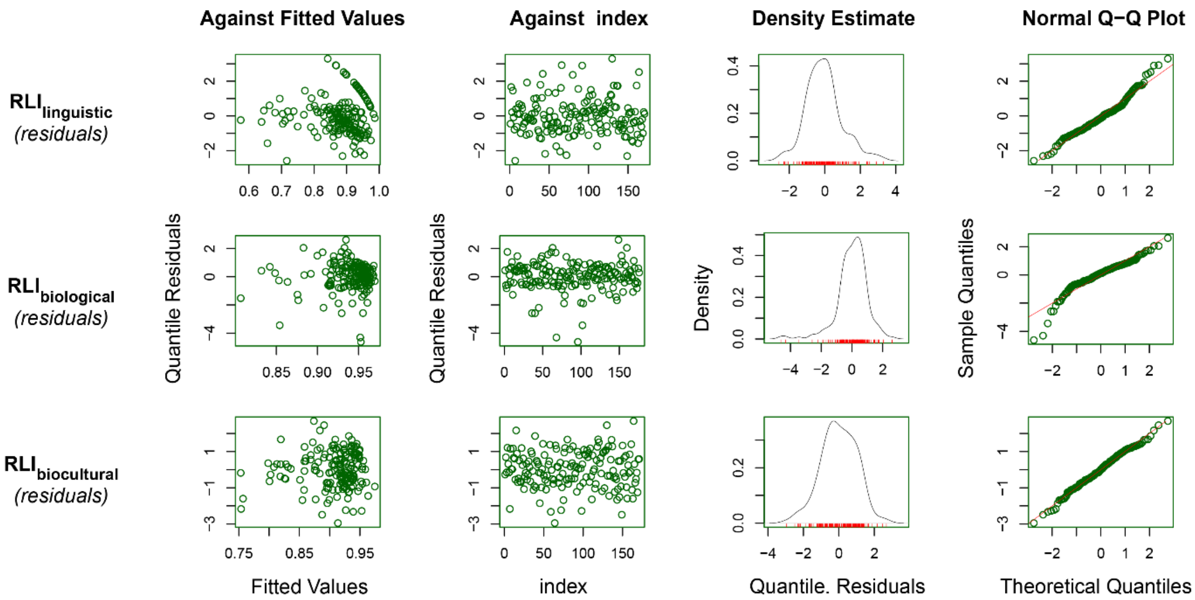

**Fig. S8.** Diagnostic plots of the hotspot models for (A) linguistic, (B) biological, and (C) biocultural diversity.

**Table S2.** Model results for the hotspot analysis using generalized linear models analyzing the effect of area and sampling bias on the different threat indicators: A) $RLI_{linguistic}$ = linguistic diversity, B) $RLI_{biological}$ = biological diversity, C) $RLI_{biocultural}$ = biocultural diversity, D) $RLI_{amphibians}$, E) $RLI_{birds}$, F) $RLI_{mammals}$, and G) $RLI_{reptiles}$. Area and sampling bias were both log-transformed and subsequently scaled.

A) Linguistic diversity

| **Predictor** | **Estimate** | **Std. Error** | **p – value** |
|---|---|---|---|
| ***Intercept*** | **2.10** | **0.10** | **< 0 .001** |
| ***Area*** | **-0.57** | **0.08** | **< 0.001** |

B) Biological diversity

| **Predictor** | **Estimate** | **Std. Error** | **p – value** |
|---|---|---|---|
| ***Intercept*** | **2.77** | **0.05** | **< 0.001** |
| *Area* | 0.03 | 0.05 | 0.51 |
| ***Sampling bias*** | **0.19** | **0.05** | **< 0.001** |
| *Area x sampling bias* | -0.04 | 0.03 | 0.22 |

C) Biocultural diversity

| **Predictor** | **Estimate** | **Std. Error** | **p – value** |
|---|---|---|---|
| ***Intercept*** | **2.30** | **0.06** | **< 0.001** |
| ***Area*** | **-0.21** | **0.06** | **< 0.001** |

D) Biological diversity - amphibans

| **Predictor** | **Estimate** | **Std. Error** | **p – value** |
|---|---|---|---|
| ***Intercept*** | **2.50** | **0.13** | **< 0.001** |
| *Area* | -0.16 | 0.09 | 0.072 |
| *Sampling bias* | 0.06 | 0.09 | 0.543 |
| *Area x sampling bias* | -0.12 | 0.07 | 0.092 |

E) Biological diversity - birds

| **Predictor** | **Estimate** | **Std. Error** | **p – value** |
|---|---|---|---|
| ***Intercept*** | **3.20** | **0.05** | **< 0.001** |
| *Area* | -0.03 | 0.05 | 0.603 |
| ***Sampling bias*** | **0.21** | **0.06** | **< 0.001** |
| *Area x sampling bias* | -0.05 | 0.03 | 0.115 |

F) Biological diversity - mammals

| Predictor | Estimate | Std. Error | p – value |
|---|---|---|---|
| ***Intercept*** | **2.46** | **0.04** | **< 0.001** |
| *Area* | -0.02 | 0.04 | 0.673 |
| ***Sampling bias*** | **0.14** | **0.04** | **< 0.001** |
| *Area x sampling bias* | -0.01 | 0.03 | 0.697 |

G) Biological diversity - reptiles

| Predictor | Estimate | Std. Error | p – value |
|---|---|---|---|
| ***Intercept*** | **2.48** | **0.08** | **< 0.001** |
| *Area* | 0.19 | 0.08 | 0.016 |
| ***Sampling bias*** | **0.47** | **0.08** | **< 0.001** |
| *Area x sampling bias* | 0.03 | 0.05 | 0.512 |

**Table S3.** Hotspot regions of linguistic, biological and biocultural diversity threat

A) Linguistic diversity

| Country | $RLI_{linguistic}$ | $RLI_{linguistic}$ (residuals) |
|---|---|---|
| Australia | *0.24* | *-2.48* |
| Taiwan | *0.52* | *-2.41* |
| El Salvador | *0.60* | *-2.10* |
| Israel | *0.65* | *-1.89* |
| Nicaragua | *0.58* | *-1.86* |

B) Biological diversity

| Country | $RLI_{biological}$ | $RLI_{biological}$ (residuals) |
|---|---|---|
| Mauritius | *0.68* | *-3.97* |
| Haiti | *0.77* | *-3.52* |
| Madagascar | *0.76* | *-3.46* |
| New Zealand | *0.73* | *-3.02* |
| Dominican Republic | *0.82* | *-2.83* |

C) Biocultural diversity

| Country | $RLI_{biocultural}$ | $RLI_{biocultural}$ (residuals) |
|---|---|---|
| Australia | *0.58* | *-3.05* |
| United States of America | *0.63* | *-2.57* |
| Haiti | *0.72* | *-2.53* |
| Taiwan | *0.72* | *-2.49* |
| Japan | *0.74* | *-2.00* |

**Table S4.** Coldspot regions of linguistic, biological and biocultural diversity threat

A) Linguistic diversity

| **Country** | $RLI_{linguistic}$ | $RLI_{linguistic}$ ***(residuals)*** |
|---|---|---|
| Saudi Arabia | *0.88* | *3.46* |
| Madagascar | *0.85* | *2.88* |
| Bulgaria | *1.00* | *2.14* |
| Uruguay | *1.00* | *2.33* |
| Democratic People’s Republic of Korea | *1.00* | *2.18* |

B) Biological diversity

| **Country** | $RLI_{biological}$ | $RLI_{biological}$ ***(residuals)*** |
|---|---|---|
| Grenada | *0.95* | *2.65* |
| Suriname | *0.99* | *1.94* |
| Trinidad and Tobago | *0.98* | *1.28* |
| Kyrgyzstan | *0.97* | *1.27* |
| Guyana | *0.98* | *1.21* |

C) Biocultural diversity

| **Country** | $RLI_{biocultural}$ | $RLI_{biocultural}$ ***(residuals)*** |
|---|---|---|
| Saudi Arabia | *0.98* | *1.99* |
| Czechia | *0.99* | *1.81* |
| Slovakia | *0.99* | *1.67* |
| Niger | *0.97* | *1.65* |
| Mali | *0.97* | *1.61* |

**Residual analysis**

**Table S5.** Model output for the driver analysis for the threats level of (A) linguistic, (B) biological diversity, and (C) biocultural.

*A) Linguistic diversity*

| ***Predictors*** | ***Estimates*** | ***Std. error*** | ***T value*** | ***P – value*** |
|---|---|---|---|---|
| ***Intercept*** | **2.16** | **0.09** | **23.37** | **<0.001** |
| ***Area*** | **-0.64** | **0.09** | **-7.04** | **<0.001** |
| *Roughness* | -0.10 | 0.07 | -1.33 | 0.185 |
| *Aridity Index* | -0.14 | 0.08 | -1.80 | 0.073 |
| *Remoteness* | -0.04 | 0.13 | -0.29 | 0.771 |
| ***Prop. urban area*** | **-0.17** | **0.06** | **-2.69** | **0.008** |
| *Prop. intense agriculture* | 0.06 | 0.09 | 0.72 | 0.474 |
| *Prop. mixed agriculture* | -0.03 | 0.10 | -0.31 | 0.757 |
| *Years of schooling* | -0.12 | 0.08 | -1.47 | 0.142 |
| ***Occupation time*** | **-0.21** | **0.07** | **-2.89** | **0.004** |

*B) Biological diversity*

| ***Predictors*** | ***Estimates*** | ***Std. error*** | ***T value*** | ***P – value*** |
|---|---|---|---|---|
| ***Intercept*** | **2.74** | **0.04** | **69.04** | **<0.001** |
| *Area* | 0.06 | 0.05 | 1.24 | 0.218 |
| ***Sampling effort*** | **0.21** | **0.04** | **4.83** | **<0.001** |
| *Area x sampling effort* | -0.01 | 0.03 | -0.36 | 0.722 |
| ***Roughness*** | **-0.10** | **0.03** | **-2.83** | **0.006** |
| ***Aridity Index*** | **-0.07** | **0.04** | **-1.85** | **0.067** |
| *Remoteness* | -0.05 | 0.05 | -0.98 | 0.327 |
| *Prop. urban area* | -0.02 | 0.03 | -0.56 | 0.576 |
| ***Prop. intense agriculture*** | **-0.22** | **0.04** | **-4.83** | **<0.001** |
| *Prop. mixed agriculture* | -0.04 | 0.06 | -0.75 | 0.454 |
| ***GDPpc*** | **-0.16** | **0.04** | **-3.59** | **<0.001** |
| ***Occupation time*** | **-0.18** | **0.04** | **-5.09** | **0.001** |

*C) Biocultural diversity*

| ***Predictors*** | ***Estimates*** | ***Std. error*** | ***T value*** | ***P – value*** |
|---|---|---|---|---|
| ***Intercept*** | **2.27** | **0.05** | **48.32** | **<0.001** |
| ***Area*** | **-0.29** | **0.06** | **-5.07** | **<0.001** |

| | | | | |
|---|---|---|---|---|
| *Roughness* | -0.05 | 0.04 | -1.24 | 0.216 |
| ***Aridity Index*** | **-0.12** | **0.05** | **-2.59** | **0.011** |
| *Remoteness* | 0.02 | 0.09 | -0.20 | 0.839 |
| *Prop. urban area* | -0.06 | 0.04 | -1.44 | 0.152 |
| *Prop. intense agriculture* | -0.03 | 0.06 | -0.45 | 0.656 |
| *Prop. mixed agriculture* | 0.01 | 0.07 | 0.08 | 0.941 |
| *GDPpc* | -0.08 | 0.05 | -1.54 | 0.126 |
| *Years of schooling* | -0.11 | 0.06 | -1.94 | 0.054 |
| ***Occupation time*** | **-0.09** | **0.04** | **-2.05** | **0.042** |

**Table S6.** Model output for the driver analysis for the threat level of (A) amphibian, (B) bird, (C) mammal, and (D) reptile diversity.

*A) Amphibian diversity*

| ***Predictors*** | ***Estimates*** | ***Std. error*** | ***T value*** | ***P – value*** |
|---|---|---|---|---|
| ***Intercept*** | **254.66** | **0.10** | **24.41** | **< 0.001** |
| *Area* | -0.26 | 0.09 | -2.76 | 0.007 |
| *Sampling effort* | 0.13 | 0.08 | 1.63 | 0.106 |
| *Area x sampling effort* | 0.03 | 0.07 | 0.40 | 0.694 |
| ***Roughness*** | **-0.29** | **0.06** | **-4.52** | **< 0.001** |
| *Aridity Index* | -0.15 | 0.08 | -1.87 | 0.063 |
| ***Remoteness*** | **0.34** | **0.13** | **2.57** | **0.011** |
| *Prop. urban area* | -0.06 | 0.08 | -0.79 | 0.429 |
| *Prop. intense agriculture* | -0.13 | 0.09 | -1.42 | 0.158 |
| ***Prop. mixed agriculture*** | **-0.26** | **0.11** | **-2.40** | **0.018** |
| *GDPpc* | -0.13 | 0.08 | -1.54 | 0.126 |
| *Occupation time* | -0.02 | 0.08 | -0.26 | 0.793 |

*B) Bird diversity*

| ***Predictors*** | ***Estimates*** | ***Std. error*** | ***T value*** | ***P – value*** |
|---|---|---|---|---|
| ***Intercept*** | **316.97** | **0.04** | **78.92** | **< 0.001** |
| *Area* | 0.02 | 0.05 | 0.32 | 0.752 |
| ***Sampling effort*** | **0.19** | **0.04** | **4.63** | **< 0.001** |
| *Area x sampling effort* | -0.03 | 0.03 | -1.24 | 0.218 |
| *Roughness* | -0.06 | 0.04 | -1.74 | 0.085 |
| *Aridity Index* | -0.03 | 0.04 | -0.82 | 0.413 |
| ***Remoteness*** | **-0.11** | **0.05** | **-2.06** | **0.041** |
| *Prop. urban area* | -0.05 | 0.03 | -1.50 | 0.136 |
| ***Prop. intense agriculture*** | **-0.20** | **0.05** | **-4.42** | **< 0.001** |
| *Prop. mixed agriculture* | -0.03 | 0.06 | -0.51 | 0.611 |
| ***GDPpc*** | **-0.20** | **0.05** | **-4.45** | **< 0.001** |
| ***Occupation time*** | **-0.14** | **0.04** | **-3.85** | **< 0.001** |

*C) Mammal diversity*

| ***Predictors*** | ***Estimates*** | ***Std. error*** | ***T value*** | ***P – value*** |
|---|---|---|---|---|
| ***Intercept*** | **2.47** | **0.03** | **80.16** | **< 0.001** |
| *Area* | -0.01 | 0.04 | -0.21 | 0.836 |

| | | | | |
|---|---|---|---|---|
| *Sampling effort* | 0.02 | 0.04 | 0.66 | 0.512 |
| *Area x sampling effort* | -0.01 | 0.02 | -0.34 | 0.732 |
| *Roughness* | -0.02 | 0.03 | -0.60 | 0.549 |
| ***Aridity Index*** | **-0.13** | **0.03** | **-4.17** | **< 0.001** |
| *Remoteness* | -0.05 | 0.05 | -1.05 | 0.298 |
| *Prop. urban area* | -0.01 | 0.03 | -0.52 | 0.606 |
| ***Prop. intense agriculture*** | **-0.12** | **0.04** | **-3.26** | **0.001** |
| *Prop. mixed agriculture* | 0.04 | 0.05 | 0.92 | 0.360 |
| *GDPpc* | -0.04 | 0.03 | -1.20 | 0.231 |
| ***Occupation time*** | **-0.12** | **0.03** | **-4.07** | **< 0.001** |

*D) Reptile diversity*

| ***Predictors*** | ***Estimates*** | ***Std. error*** | ***T value*** | ***P – value*** |
|---|---|---|---|---|
| ***Intercept*** | **2.54** | **0.07** | **34.95** | **< 0.001** |
| ***Area*** | **0.33** | **0.08** | **3.86** | **< 0.001** |
| ***Sampling effort*** | **0.49** | **0.08** | **6.35** | **< 0.001** |
| *Area x sampling effort* | 0.09 | 0.05 | 1.75 | 0.081 |
| ***Roughness*** | **-0.17** | **0.06** | **-3.04** | **0.003** |
| *Aridity Index* | -0.03 | 0.06 | -0.43 | 0.672 |
| *Remoteness* | -0.06 | 0.08 | -0.68 | 0.500 |
| *Prop. urban area* | 0.001 | 0.05 | 0.03 | 0.978 |
| ***Prop. intense agriculture*** | **-0.25** | **0.07** | **-3.56** | **< 0.001** |
| *Prop. mixed agriculture* | 0.03 | 0.09 | 0.32 | 0.751 |
| *GDPpc* | 0.02 | 0.07 | 0.28 | 0.781 |
| ***Occupation time*** | **-0.34** | **0.06** | **-6.15** | **< 0.001** |

**Table S7.** Model output for the driver analysis for the threat level of (A) biocultural, and (B) linguistic diversity including GDPpc instead of schooling.

*A) Biocultural diversity*

| ***Predictors*** | ***Estimates*** | ***Std. error*** | ***T value*** | ***P – value*** |
|---|---|---|---|---|
| ***Intercept*** | **225.16** | **0.05** | **47.74** | **< 0.001** |
| ***Area*** | **-0.29** | **0.06** | **-5.07** | **< 0.001** |
| *Roughness* | -0.05 | 0.04 | -1.28 | 0.204 |
| ***Aridity Index*** | **-0.13** | **0.05** | **-2.86** | **0.005** |
| *Remoteness* | 0.03 | 0.09 | 0.35 | 0.731 |
| *Prop. urban area* | -0.06 | 0.04 | -1.51 | 0.134 |
| *Prop. intense agriculture* | -0.03 | 0.06 | -0.44 | 0.664 |
| *Prop. mixed agriculture* | 0.01 | 0.07 | 0.21 | 0.834 |
| ***GDPpc*** | **-0.11** | **0.05** | **-2.36** | **0.019** |
| *Occupation time* | -0.08 | 0.04 | -1.87 | 0.063 |

*B) Linguistic diversity*

| ***Predictors*** | ***Estimates*** | ***Std. error*** | ***T value*** | ***P – value*** |
|---|---|---|---|---|
| ***Intercept*** | **215.45** | **0.09** | **23.46** | **< 0.001** |
| ***Area*** | **-0.62** | **0.09** | **-6.85** | **< 0.001** |
| *Roughness* | -0.11 | 0.07 | -1.51 | 0.134 |
| ***Aridity Index*** | **-0.17** | **0.08** | **-2.20** | **0.029** |
| *Remoteness* | -0.07 | 0.14 | -0.51 | 0.614 |
| ***Prop. urban area*** | **-0.17** | **0.06** | **-2.64** | **0.009** |
| *Prop. intense agriculture* | 0.04 | 0.09 | 0.478 | 0.635 |
| *Prop. mixed agriculture* | -0.06 | 0.10 | -0.61 | 0.541 |
| *GDPpc* | -0.14 | 0.08 | -1.70 | 0.092 |
| ***Occupation time*** | **-0.18** | **0.07** | **-2.49** | **0.014** |

**Sensitivity analysis**

To test the robustness of our results for the driver analysis we additionally ran models using the residuals of the hotspot analysis as response variable. Using the residuals of the hotspot models follows the assumption that they are independent of the effect of area for biocultural and linguistic diversity and of the effect of area and sampling bias in the case of the biological diversity. Hence, we build linear models with a Gaussian error distribution including all driver variables as in the other model excluding area and sampling effort. Model fit was checked visually and autocorrelation was checked using Durbin-Watson tests that indicated not correlation in the model residuals (see table S6).

**Table S8.** Model results of the sensitivity analysis for the driver models. Here we used the residuals of the hotspot analysis as response variables for (A) linguistic diversity, (B) biological diversity, and (C) biocultural diversity.

*A) Linguistic diversity*

| ***Predictors*** | ***Estimates*** | ***Std. error*** | ***T value*** | ***P – value*** |
|---|---|---|---|---|
| *Intercept* | -0.03 | 0.07 | -0.44 | 0.662 |
| *Roughness* | -0.11 | 0.08 | -1.48 | 0.141 |
| *Aridity Index* | -0.16 | 0.08 | -1.89 | 0.061 |
| *Remoteness* | -0.08 | 0.13 | -0.58 | 0.566 |
| *Prop. urban area* | -0.12 | 0.08 | -1.54 | 0.127 |
| *Prop. intense agriculture* | 0.01 | 0.10 | 0.15 | 0.884 |
| *Prop. mixed agriculture* | -0.06 | 0.11 | -0.55 | 0.585 |
| *GDPpc* | -0.18 | 0.12 | -1.43 | 0.156 |
| *Years of schooling* | -0.00 | 0.13 | -0.01 | 0.993 |
| ***Occupation time*** | **-0.17** | **0.08** | **-2.07** | **0.040** |

*B) Biological diversity*

| ***Predictors*** | ***Estimates*** | ***Std. error*** | ***T value*** | ***P – value*** |
|---|---|---|---|---|
| *Intercept* | 0.02 | 0.07 | 0.24 | 0.809 |
| *Roughness* | -0.12 | 0.07 | -1.71 | 0.089 |
| ***Aridity Index*** | **-0.24** | **0.08** | **-3.14** | **0.002** |
| *Remoteness* | -0.10 | 0.10 | -0.95 | 0.344 |
| *Prop. urban area* | -0.02 | 0.07 | 0.00 | 0.753 |
| ***Prop. intense agriculture*** | **-0.31** | **0.08** | **-3.79** | **<0.001** |
| *Prop. mixed agriculture* | 0.03 | 0.09 | 0.33 | 0.738 |
| *GDPpc* | -0.17 | 0.11 | -1.56 | 0.120 |
| *Years of schooling* | 0.10 | 0.11 | 0.84 | 0.401 |

| | | | | |
|---|---|---|---|---|
| ***Occupation time*** | **-0.27** | **0.07** | **-3.76** | **<0.001** |

*C) Biocultural diversity*

| ***Predictors*** | ***Estimates*** | ***Std. error*** | ***T value*** | ***P – value*** |
|---|---|---|---|---|
| *Intercept* | 0.01 | 0.07 | 0.09 | 0.929 |
| ***Roughness*** | **-0.16** | **0.07** | **-2.38** | **0.019** |
| ***Aridity Index*** | **-0.21** | **0.08** | **-2.84** | **0.005** |
| *Remoteness* | -0.12 | 0.12 | -0.99 | 0.325 |
| *Prop. urban area* | -0.07 | 0.07 | -0.90 | 0.371 |
| *Prop. intense agriculture* | -0.11 | 0.09 | -1.19 | 0.235 |
| *Prop. mixed agriculture* | 0.08 | 0.10 | 0.76 | 0.452 |
| *GDPpc* | -0.13 | 0.09 | -1.46 | 0.147 |
| *Years of schooling* | -0.13 | 0.09 | -0.46 | 0.146 |
| ***Occupation time*** | **-0.31** | **0.07** | **-4.21** | **<0.001** |

**Table S9.** Model fit for the driver models (cf. Tables S5 and S6) across diversity threat variables.

| ***Diversity threat variable*** | ***R squared*** |
|---|---|
| $RLI_{linguistic}$ | **0.37** |
| $RLI_{biological}$ | **0.42** |
| $RLI_{biocultural}$ | **0.40** |
| $RLI_{amphibians}$ | **0.49** |
| $RLI_{birds}$ | **0.43** |
| $RLI_{mammals}$ | **0.34** |
| $RLI_{reptiles}$ | **0.45** |
